# Sleep deprivation selectively down-regulates astrocytic 5-HT_2B_ receptors and triggers depressive-like behaviors via stimulating P2X_7_ receptors

**DOI:** 10.1101/745505

**Authors:** Maosheng Xia, Shuai Li, Shanshan Liang, Xiaowei Li, Zexiong Li, Alexei Verkhratsky, Dawei Guan, Baoman Li

## Abstract

Chronic loss of sleep damages health and disturbs quality of life. The long-lasting sleep deprivation (SD) as well as sleep abnormalities is a substantial risk factor for major depressive disorder (MDD), although the underlying mechanisms are not clear. In our previous studies, we report the activation of nucleotide-binding domain and leucine-rich repeat protein-3 (NLRP3) inflammasome induced by long-term SD is P2X_7_ receptors (P2X_7_R) dependent, and antidepressant fluoxetine could alleviate this neuroinflammasome via 5-HT_2B_ receptors (5-HT_2B_R) in astrocytes. Here, we discovered that the chronic SD activates astroglial P2X_7_ receptors, which in turn selectively down-regulated expression of 5-HT_2B_R in astrocytes. Stimulation of P2X_7_R induced by SD suppressed the phosphorylation of AKT and FoxO3a selectively in astrocytes, but not in neurones. The over-expression of FoxO3a in astrocytes inhibited expression of 5-HT_2B_R. Down-regulation of 5-HT_2B_R instigated by SD suppressed activation of STAT3 and relieved the inhibition of Ca^2+^-dependent phospholipase A2 (cPLA2). This latter cascade promoted the release of arachidonic acid (AA) and prostaglandin E2 (PGE2). The depressive-like behaviours induced by SD were alleviated in P2X_7_R-KO mice. Our study reveals the mechanism underlying chronic SD-induced depressive-like behaviors and highlights that blocking P2X_7_ receptors or activating 5-HT_2B_R in astrocytes could play a key role for exploring the therapeutic strategies aimed at the depression evoked by sleep disorders.

**Main Points:** Chronic SD selectively down-regulates expression of 5-HT_2B_R through activation of P2X_7_R in astrocytes. SD promotes the release of AA and PGE2 via the decreased 5-HT_2B_R, these factors induce depressive-like behaviors.

## 1. Introduction

Good sleep safeguards physical health and quality of life, and contributes to cognitive and emotional functions. Prolonged sleep deprivation (SD) increases the risk of mood disorders (Alvaro, Roberts, Harris, & Bruni, 2017) and impairs regulation of emotions (Goldstein & Walker, 2014). Our previous studies revealed that chronic SD induces depressive-like behaviours via stimulating an astroglial neuroinflammatory response. This response is linked to activation of P2X_7_ purinoceptors associated with the activation of nucleotide-binding domain and leucine-rich repeat protein-3 (NLRP3) inflammasome. We also found that the anti-depressant fluoxetine suppresses the activation of NLRP3 inflammasome caused by SD via astroglial 5-HT_2B_ receptors (Li et al., 2018; Xia et al., 2018). However, the details P2X_7_R and 5-HT_2B_R interactions are still unknown, especially in the context of chronic SD.

Dynamic changes of brain extracellular ATP during sleep-wake cycle have not been studied extensively. Increased ATP release from neurones during the period of wakefulness was nonetheless suggested to activate P2X_7_R in neural cells (Chennaoui et al., 2017). Functional P2X_7_Rs are distributed in cortex, hippocampus and retina, and P2X_7_R interferes with the sleep rhythms directly or indirectly through the release of cytokines and neurotransmitters (Illes, Verkhratsky, Burnstock, & Franke, 2012). Expression of P2X_7_Rs was up-regulated in people subjected to chronic SD; and these changes in expression were linked to the cycling in bipolar disorder (Backlund et al., 2012).

The 5-HT_2B_Rs are related to the major depressive disorder (MDD); these receptors expressed in astrocytes are targets for serotonin-specific re-uptake inhibitors (SSRIs) (Li, Zhang, Li, Hertz, & Peng, 2009; Li, Zhang, Zhang, Hertz, & Peng, 2011; Li et al., 2008; Li et al., 2018; Peng, Gu, Li, & Hertz, 2014; Peng, Song, Li, & Verkhratsky, 2018). Down-regulation of 5-HT_2B_Rs leads to a loss of sleep homeostasis in Drosophila (Qian et al., 2017). We found that leptin enhances the anti-depressive potential of fluoxetine in the context of SD-induced depression by stimulating 5-HT_2B_R in astrocytes with consequent increase in the phosphorylation of ERK_1/2_ (Li, Zhang, Li, Hertz, & Peng, 2010). We also found that 5-HT_2B_Rs regulate expression of Ca^2+^-dependent phospholipase A2 (cPLA2) via transactivation of epidermal growth factor receptor (EGFR) (Li et al., 2009). In spinal cord astrocytes, the phosphorylation of cPLA2 induced by ATP induces the rapid release of arachidonic acid (AA) and prostaglandin E2 (PGE2) (Xia & Zhu, 2011).

Ionotropic P2X_7_Rs promote NLRP3 inflammasome assembly and trigger ATP-induced release of mature IL-1β and IL-18 in astrocytes (Xia et al., 2018). Activation of P2X_7_R is linked to NLRP3 inflammasome and induction of depressive-like behaviours induced by chronic stress (Yue et al., 2017). Furthermore, P2X_7_Rs suppress phosphorylation of AKT and ERK induced by BzATP in microglia (Kim, Ko, Hyun, Min, & Kang, 2018). We demonstrated that SD could decrease the phosphorylation of AKT, but not the phosphorylation of ERK (Xia et al., 2018), while activation of AKT can phosphorylate the Forkhead transcriptional factor FoxO3a on Ser253 site; the FoxO3a is an attractive candidate for regulating stress responses (Polter et al., 2009).

In this study, we dissect how SD-induced activation of P2X_7_Rs regulates expression of 5-HT_2B_R and the production of arachidonic acid (AA) and prostaglandin E2 (PGE2) to reveal the possible mechanism linking chronic SD with mood disorders including the MDD.

## 2. Materials and Methods

### 2.1 Animals

As in our previous studies (Xia et al., 2018; Xia, Yang, Sun, Qi, & Li, 2017), C57BL/6, B6.Cg-Tg(Thy1-YFP)HJrs/J, FVB/N-T g(GFAP-GFP) 14Mes/J and B6.129P2-P2rx7^tm1Gab^/J (P2X_7_R-KO) mice were all purchased from the Jackson Laboratory (Bar Harbor, ME, USA). Male mice were used at an age of approximately 3 months (~25 g) and were kept in standard housing conditions (22 ± 1°C; light/dark cycle of 12 h/12 h) with food and water available *ad libitum*. All mice were randomly assigned to different experimental groups with a random number table. All operations were carried out in accordance with the USA National Institutes of Health Guide for the Care and Use of Laboratory Animals (NIH Publication No. 8023) and its 1978 revision, and all experimental protocols were approved by the Institutional Animal Care and Use Committee of China Medical University, No. [2019]059.

### 2.2 Materials

Most chemicals, including BzATP (2’(3’)-O-(4-Benzoylbenzoyl)adenosine 5’-triphosphate triethylammonium salt), DAPI (4’,6-diamidine-2-phenylindole) dihydrochloride), BW723C86 (α-methyl-5-(2-thienylmethoxy)-1H-Indole-3-ethanamine monohydrochloride) and a primary antibody raised against β-actin were purchased from Sigma (USA). Other primary antibodies, Alexa Fluor-conjugated or horseradish peroxidase-conjugated secondary antibodies were purchased from Millipore (USA). Stattic (6-Nitrobenzo[b]thiophene 1,1-dioxide) was supplied by R&D Systems (UK).

Chemicals for the preparation of cell sorting medium were obtained from Gibco Life Technology Invitrogen (USA).

### 2.3 Sleep Deprivation (SD)

As described previously (Franken, Dijk, Tobler, & Borbely, 1991), the SD was maintained by “gentle handling” according to the standard protocols, gently touching the mouse with a brush to keep it awake in every cage. SD was operated for 6 hours, which began at 7 a.m. and ended at 1 p.m. During SD, the mice continually were offered food and water *ad libitum*. Animals in the sham group were preserved undisturbed in a separate room with the same light/dark rhythm as SD group. The mice were treated with sham or SD stimulation for 3 or 4 weeks.

### 2.4 Tissue Collection and Measurement of ATP-Content

Tissues were collected as described before (Dworak, McCarley, Kim, Kalinchuk, & Basheer, 2010). Mice were killed by decapitation. Brain slices (2 mm thick) were carefully placed on dry ice containing covered Petri-dish for the rapid freezing and the ensuing dissection, frontal cortex region was selected. Extreme care was observed to complete this process rapidly; all the tissue samples were collected at 10 a.m. during the light period at the end of every week. The selected regions were held frozen on dry ice and stored at −80° C until used for the left measurements.

ATP levels were assessed with luciferin-luciferase based assay (Lundin & Thore, 1975; McElroy & DeLuca, 1983) using a commercial ATP assay system with bioluminescence detection kit (Enliten, Promega). ATP was measured according to the manufacturer’s protocol. In brief, weighed tissue samples were homogenised in 5% trichloroacetic acid (TCA) and centrifuged at 5000 rpm in cold for 5 min, the supernatant was transferred to a fresh tube. Samples were neutralised with Tris acetate buffer, the luciferase reagent was used immediately before measurement in the luminometer. The measured concentration was normalised to “0 week” point of sham group.

### 2.5 Cells Dissociation and Fluorescence-activated Cell Sorting (FACS)

B6.Cg-Tg(Thy1-YFP)HJrs/J and FVB/N-Tg(GFAP-GFP)14Mes/J mice were separately used for isolating neurones and astrocytes. A single-cell suspension from the cortex and hippocampus was prepared as previously described (Xia et al., 2017). In brief, tissue from 3 mice was pooled for one sample. Wavelengths of 488 nm and 530/30 nm were used for YFP or GFP excitation and emission, respectively. YFP^+^ or GFP^+^ cells were sorted and collected. As this method, the purity of neurones or astrocytes sorted has been identified in our previous reports (Fu, Li, Hertz, & Peng, 2012) by checking mRNA level of cell specific markers of astrocytes, neurons and oligodendrocytes.

### 2.6 Primary Culture of Astrocytes

As described previously (Li et al., 2011; Li et al., 2008; Xia et al., 2017), astrocytes isolated from the cerebral hemispheres of newborn C57BL/6 mice were cultured in Dulbecco’s modified Eagle’s medium (DMEM) with 7.5 mM glucose. From the third week, dibutyryl cyclic AMP (dBcAMP) was added to the medium.

### 2.7 Immunohistochemistry

The brain tissue was fixed and immersed in 4% paraformaldehyde (PFA) and cut in 100 μm slices. Immunohistochemistry was performed as previously described (Li et al., 2016; Xia et al., 2017). In brief, the following primary antibodies were used: mouse anti-5-HT_2B_R (1:150), mouse anti-P2X_7_ (1:150), chicken anti-NeuN (1:250), rabbit anti-Glt1 (1:200), rabbit anti-GFAP (1:250). The slices were incubated with Alexa Fluor-conjugated secondary antibodies for 2 hours at room temperature (1:250). DAPI (1:2000) was stained to identify cell nuclei. Immunofluorescence was imaged using a confocal scanning microscope (DMi8, Leica, Germany). The background intensity of each image was calculated in cell-free parenchyma in the same field of view and subtracted from the total immunofluorescence intensity. The intensity of 5-HT_2B_R immunofluorescence from each group was normalised to the intensity of the sham group.

### 2.8 Western Blotting

As described previously (Li et al., 2016; Li et al., 2011), the sections were blocked with powdered skim milk and incubated for 2 h with the primary antibodies at room temperature. After washing three times, specific binding was detected with horseradish peroxidase-conjugated secondary antibodies. Staining was visualised with ECL detection reagents, and images were acquired with an electrophoresis gel imaging analysis system. Band density was measured in Windows AlphaEase FC 32-bit software.

### 2.9 Real-time PCR (RT-PCR)

As described previously (Xia et al., 2017), total RNA was reverse transcribed, and PCR amplification was performed with a Thermo-cycler. The RNA quantities were normalised by glyceraldehyde 3-phosphate dehydrogenase (GAPDH) before calculating the relative expression of 5-HT_2B_R. Values were first calculated as the ratio of the relative expression of 5-HT_2B_R and GAPDH, then the values were normalised by sham group.

### 2.10 Over-expression of FoxO3a by Using Adenoviral Vectors

As described previously (Ren et al., 2018; Xia & Zhu, 2013), replication-defective adenoviral vectors expressing dominant wild type FoxO3a were designed and purchased from TaKaRa Biotechnology (Dalian, China). The wild-type FoxO3a had a hemagglutinin tag at the N-terminus and expressed green fluorescent protein (GFP). Astrocytes were infected with recombinant adenovirus in DMEM medium for 8 h, after which the medium was replaced by fresh complete culture medium including 10% foetal bovine serum. The infection efficiency was close to 80 %, as determined by the GFP expression.

### 2.11 Assessment of Arachidonic Acid (AA) Mobilisation

As described previously (Xia & Zhu, 2011, 2014), the release of ^3^H from astrocytes prelabelled with [^3^H]AA was used to monitor the response to serotonin. Confluent cultures were changed by quiescent medium for 24 h before they were labelled for 4 h with 1 μCi/ml [^3^H] arachidonic acid (PerkinElmer Life Sciences). Washing cells once with PBS containing 0.1% free fatty acid albumin and twice with PBS alone. Astrocytes were then incubated at 37 °C in fresh Ham’s F-12 medium supplemented with 0.1% free fatty acid albumin plus serotonin or PBS. The astrocytes were washed with 5% Triton and scraped off. The radioactivity levels of astrocytes and medium were quantified by scintillation counting. The results were normalised and expressed as a percentage of the mean of the basal release.

### 2.12 PGE2 Assays

As described previously (Xia & Zhu, 2011, 2014), PGE2 levels were monitored by using enzyme immunoassay kit (Cayman Chemical, Ann Arbor, MI). The assay was operated according to the manufacturer’s protocols. PGE2 production was evaluated in duplicate, and concentrations were calculated from a standard curve of PGE2 standards. The sensitivity of the assay allowed detection of up to 15 pg/ml. When necessary, the samples were diluted in the assay buffer.

### 2.13 Tail Suspension Test (TST)

TST is a behavioural despair-based test. The mouse was suspended by its tail around 2 cm from the tip as our previous description (Li et al., 2018; Xia et al., 2017), behaviour was recorded for 6 min. The duration of immobility was measured by an observer blinded to the treatment groups.

### 2.14 Forced Swimming Test (FST)

FST is also a behavioural despair-based test. In brief, each mouse was trained to swim 15 min before the formal measurement. Then the trained mouse was put into a glass cylinder that contained 30 cm deep water (25 ± 1 ° C) for 6 min. The time of immobility was recorded during the last 4 min period which followed 2 min of habituation (Li et al., 2018; Xia et al., 2017).

### 2.15 Sucrose Preference Test

As previously described (Li et al., 2018; Xia et al., 2017), the sucrose preference test is a reward-based test and a measure of anhedonia. Briefly, after 12 hours of food and water deprivation, the mice were provided with two bottles which were pre-weighed, including one bottle was filled with 2.5% sucrose solution and a second bottle contained water, for 2 h. The percentage preference was calculated according to the following formula: % preference = [sucrose intake/(sucrose + water intake)] x 100%.

### 2.16 Statistical Analysis

All measurements were performed by an investigator blinded to the experimental conditions. Differences between multiple groups were evaluated by analysis of variance (ANOVA) followed by Fisher’s least significant difference (LSD) or a Tukey-Kramer *post hoc* multiple comparison test for unequal replications using GraphPad Prism 5 software (GraphPad Software Inc., La Jolla, CA). All statistical data are expressed as the mean ± SEM; the level of significance was set at p < 0.05.

## 3. Results

### 3.1 SD Specifically Down-regulates 5-HT_2B_R expression in Astrocytes

After the treatment with SD for 3 weeks, the fluorescence intensity of 5-HT_2B_R immunoreactivity was significantly decreased by 68 ± 7.1% (n = 6) in astrocytes when compared with a sham group; there was no obvious difference in 5-HT_2B_Rs expression in neurones between sham and SD groups (Fig. 1A, B). We further measured expression of 5-HT_2B_R in neurones and astrocytes FACS sorted from Thy1-YFP and GFAP-GFP mice treated with SD. As shown in Fig. 1C, the mRNA level of 5-HT_2B_R was significantly suppressed by 71 ± 5.2% (n = 6) in astrocytes, however, SD marginally and nonsignificantly affected expression of 5-HT_2B_R in neurones.

**Figure 1.**
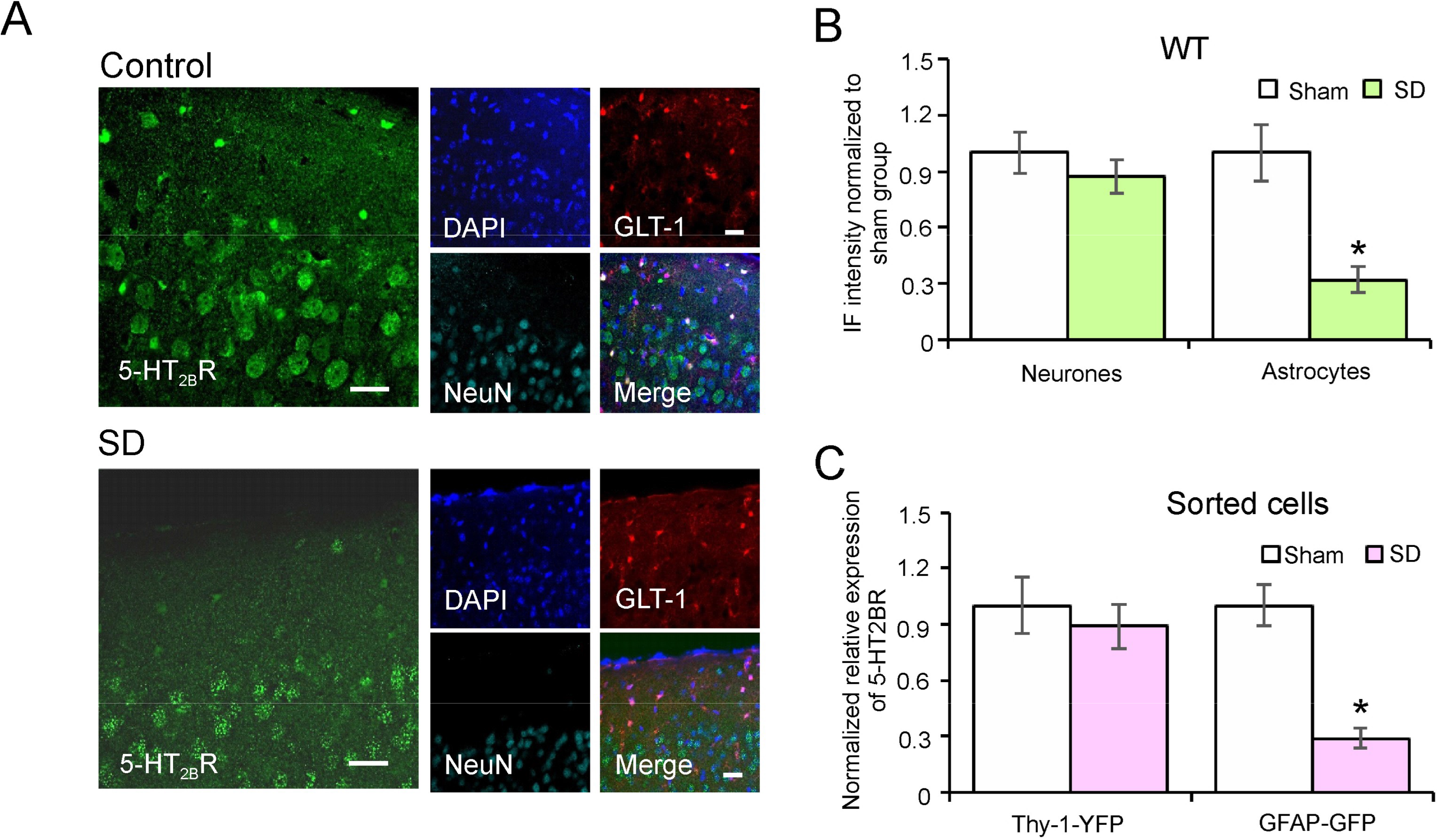
5-HT_2B_ receptors was selectively decreased by the treatment with SD in astrocytes. (A) Immunolabelled 5-HT_2B_ receptors (green) were co-stained with DAPI (blue), Glt1 (red) and NeuN (cyan) in the mice cortex treated with sham (Control) or exposed to SD for 3 weeks. Scale bar, 20 μm. (B) 5-HT_2B_ receptors immunolabeling intensity of neurones and astrocytes respectively relative to the cell-free parenchyma in cortex, normalised to the intensity of sham group. (C) qPCR analysis of 5-HT_2B_ receptors mRNA expression in FACS sorted cortical neurons and astrocytes, expressed as the relative expression ratio normalised to sham group. Data represent mean ± SEM. *Indicates statistically significant (p<0.05) difference from sham group, n = 6.

### 3.2 SD Induced Downregulation of 5-HT_2B_R is Mediated by P2X_7_R

The SD induced down-regulation of 5-HT_2B_Rs was completely eliminated in astrocytes from P2X_7_R-KO mice (expression of P2X_7_R in P2X_7_R-KO mice was significantly suppressed, as shown in Fig 2A). The fluorescence intensity of 5-HT_2B_R immunoreactivity in astrocytes was 92 ± 15.2% of control group, whereas it was 105 ± 10.8% of control group in neurones (Fig. 2B, C).

**Figure 2.**
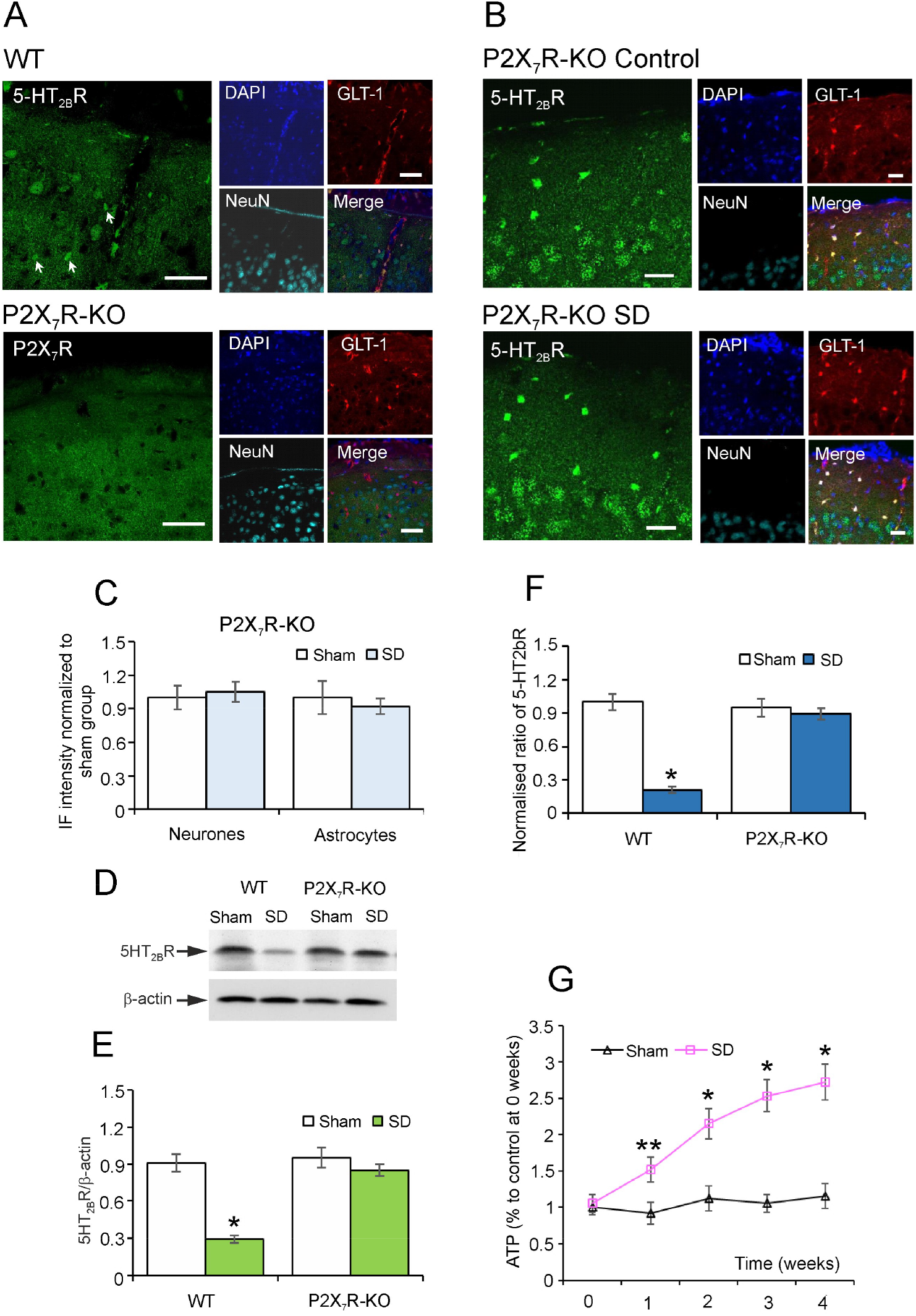
P2X_7_ receptors mediate SD-dependent decrease in expression of 5-HT_2B_ receptors. (A) Immunolabelled of P2X_7_R (green) were co-stained with DAPI (blue), Glt1 (red) and NeuN (cyan) in the cortex of wild type (WT) and P2X_7_R-KO mice. Scale bar, 50 μm. (B) Immunolabelled of 5-HT_2B_ receptors (green) were co-stained with DAPI (blue), Glt1 (red) and NeuN (cyan) in the cortex of P2X_7_R-KO mice treated with sham (Control) or exposed to SD for 3 weeks. Scale bar, 20 μm. (C) 5-HT_2B_ receptors immunolabelling intensity of neurons and astrocytes respectively relative to the cell-free parenchyma in cortex, normalised to the intensity of sham group. (D) The protein expression of 5-HT_2B_ receptors in wild type or P2X_7_R-KO mice treated with sham (Control) or exposed to SD for 3 weeks was calculated as the ratio of 5-HT_2B_R and β-actin (E). Data represent mean ± SEM. *Indicates statistically significant (p<0.05) difference from sham group, n = 6. (F) qPCR analysis of 5-HT_2B_ receptors mRNA expression in wild type or P2X_7_R-KO mice treated with or without SD for 3 weeks, expressed as the relative expression ratio normalised to sham group. Data represent mean ± SEM. *Indicates statistically significant (p<0.05) difference from sham group, n = 6. (G) The level of ATP in the cortex of wild type mice treated with SD from 0 to 4 weeks, normalised to sham group at 0 week. Data represent mean ± SEM. *Indicates statistically significant (p<0.05) difference from sham group at the same time point; **indicates statistically significant (p<0.05) difference from any other group, n = 6.

The SD-related changes in protein expression of 5-HT_2B_R quantified by western blotting, were similarly absent in P2X_7_R-KO mice (Fig. 2D). The treatment with SD significantly reduced the expression of 5-HT_2B_R to 32% of control group in wild type mice, whereas the level of 5-HT_2B_R in SD group was 89 ± 4.8% of control group in P2X_7_R-KO mice, and there was no obvious difference in the expression of 5-HT_2B_R in astrocytes from WT control and from P2X_7_R-KO mice (Fig. 2E).

The mRNA level of 5-HT_2B_R measured by RT-PCR reflected protein expression. As compared with control, mRNA expression of 5-HT_2B_R was significantly down-regulated by 79 ± 3.9% in SD group of the wild type mice (Fig. 2F). However, the treatment with SD decreased expression of 5-HT_2B_R mRNA only by 6 ± 2.5% of control in P2X_7_R-KO mice (Fig. 2F).

Meanwhile, we measured the effect of 4 weeks of SD on ATP levels in the frontal cortex. As shown in Fig. 2G, treatment with SD resulted in gradual increase of the level of ATP. As compared with the initial point (0 week) and control groups SD significantly increased ATP by 52 ± 17.1% at 1 week, by 107 ± 20.7% at 2 weeks, by 143 ± 22.2% at 3 weeks, by 151 ± 25.4% at 4 weeks (n = 6); there were no statistical difference in the level of ATP from 2 to 4 weeks. (Fig. 2G). Moreover, there was no significant fluctuation of ATP level from 0 to 4 weeks in control group (Fig. 2G).

### 3.3 The Involvement of AKT and FoxO3a in Regulation of 5-HT_2B_R by P2X_7_R *in vivo* and *in vitro*

In wild type mice treated with SD, the phosphorylation of AKT (Ser473) was significantly reduced by 58 ± 3.3% (n = 6) of control group (Fig. 3A), the phosphorylation of FoxO3a (Ser253) was decreased by 61 ± 5.4% (n = 6) as compared with controls (Fig. 3B). To the contrary in P2X_7_R-KO mice treated with SD, the phosphorylation of AKT was somewhat increased to 116 ± 16.4% (n = 6) of control group (Fig. 3A), whereas the level of p-FoxO3a was 123 ± 12.3% (n = 6) of control group (Fig. 3B); there were no significant difference between control and SD groups.

**Figure 3.**
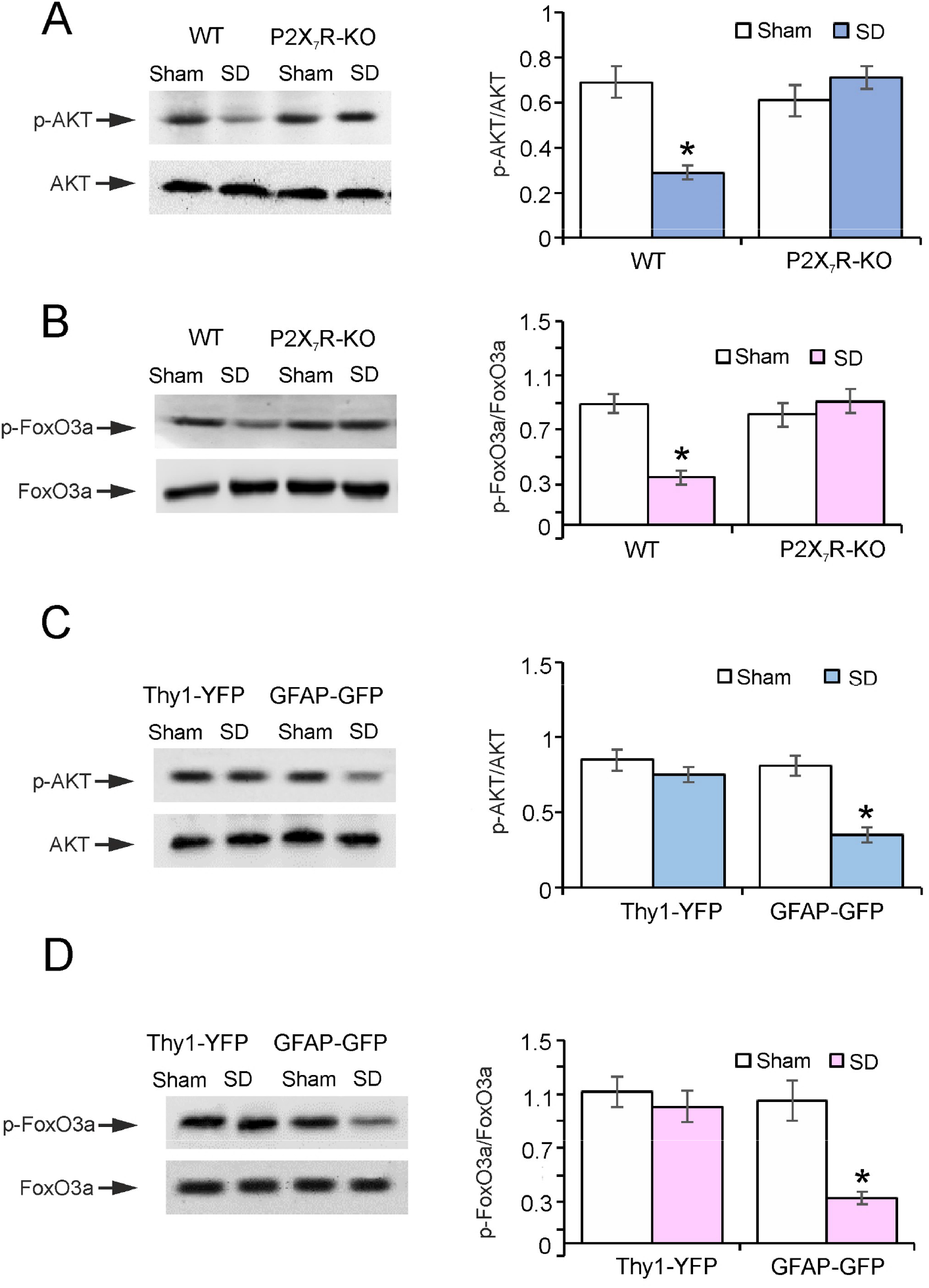
Signalling cascades involved in regulation of expression of 5HT_2B_R. (A and B) The wild type (WT) and P2X_7_R-KO mice were treated with sham (Control) or exposed to SD for 3 weeks, the ratio of p-AKT and AKT (A), and the ratio of p-FoxO3a and FoxO3a (B) were analyzed. (C and D) Thy1-YFP and GFAP-GFP mice were treated with or without SD for 3 weeks, the sorted neurones and astrocytes were collected to measure the level of p-AKT/AKT (C) and p-FoxO3a/FoxO3a (D). Data represent mean ± SEM. *Indicates statistically significant (p<0.05) difference from sham group, n=6.

We used SD-treated Thy1-YFP or GFAP-GFP transgenic mice to measure the levels of p-AKT and p-FoxO3a in the FACS sorted neurones or astrocytes. As shown in Fig. 3C, the treatment with SD selectively decreased the phosphorylation of AKT in astrocytes by 57 ± 4.7% (n = 6) of control group, whereas no obvious difference existed between control and SD groups in neurones. Similarly, the SD suppressed the phosphorylation of FoxO3a by 69 ± 5.2% (n = 6) in astrocytes as compared with controls, while no changes in neurones were observed (Fig. 3D).

In primary cultured astrocytes, we used P2X_7_R agonist (BzATP) to simulate the effects of ATP on P2X_7_R induced by SD. For probing the regulation mechanism of P2X_7_R on the expression of 5-HT_2B_R in the primary cultured astrocytes, mRNA of P2X_7_R was RNA interfered with siRNA duplex. This manipulation decreased the level of P2X_7_R by 88 ± 3.4% of control group (Fig. 4A). For identifying the effect of transcription factor FoxO3a on the expression of 5-HT_2B_R, we over-expressed the FoxO3a in primary cultured astrocytes, as shown in Fig. 4B.

**Figure 4.**
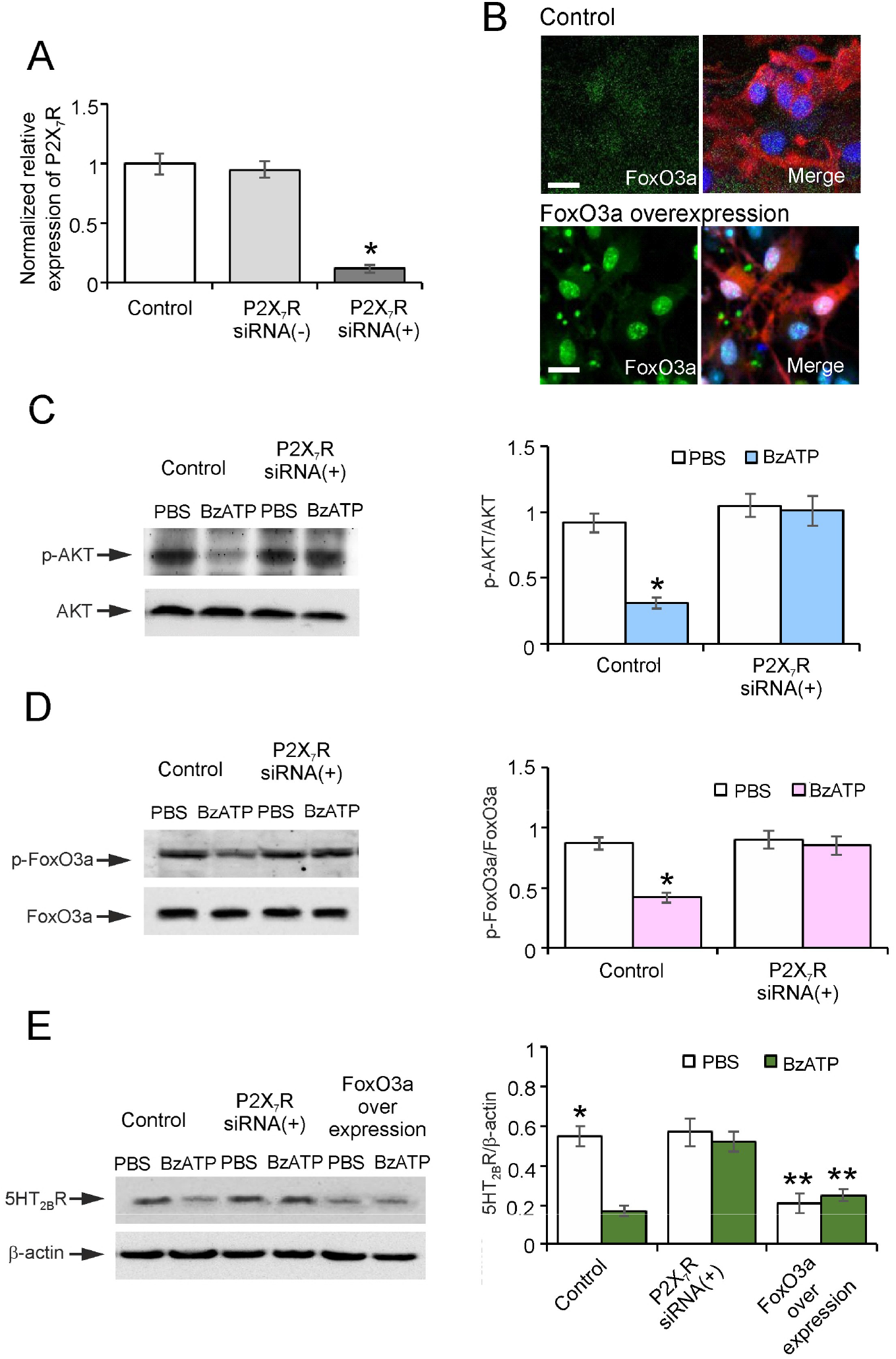
Role of P2X_7_ receptors in regulation of the expression of 5-HT_2B_ receptors *in vivo*. (A) qPCR analysis of P2X_7_R mRNA expression in the primary cultured astrocytes treated with negative control or P2X_7_R siRNA duplex for 3 days, expressed as the relative expression ratio normalised to control group. Data represent mean ± SEM. *Indicates statistically significant (p<0.05) difference from control group, n = 6. (B) Immunohistochemistry of FoxO3a (green) were co-stained with DAPI (blue) and GFAP (red) in astrocytes infected with or without recombinant adenovirus for 3 days. Scale bar, 20 μm. (C and D) The primary cultured astrocytes were pre-treated with P2X_7_R siRNA duplex for 3 days, the ratio of p-AKT and AKT (C2) and the ratio of p-FoxO3a and FoxO3a (D2) were checked. Data represent mean ± SEM. *Indicates statistically significant (p<0.05) difference from control group, n = 6. (E) The primary cultured astrocytes were pre-treated with P2X_7_R siRNA duplex or over-expressed FoxO3a with recombinant adenovirus for 3 days, the protein expression was expressed as the ratio of 5-HT_2B_R and β-actin (E2). Data represent mean ± SEM. *Indicates statistically significant (p<0.05) difference from PBS group; **indicates statistically significant (p<0.05) difference from any other group except each other, n = 6.

Administration of BzATP reduced the phosphorylation of AKT by 66 ± 4.3% of PBS control group in astrocytes (Fig. 4C). In contrast, after treatment with siRNA interfering, the effect of BzATP on the phosphorylation of AKT was abolished, as there was no significant difference between PBS and BzATP groups (Fig. 4C). Phosphorylation of FoxO3a was decreased by 52 ± 4.1% (n = 6) in BzATP group, as compared with PBS control (Fig. 4D). After 3 days of P2X_7_R siRNA treatment, the phosphorylation of FoxO3a treated with BzATP was recovered to 94 ± 7.7% of PBS group (Fig. 4D). In astrocytes, BzATP reduced the expression of 5-HT_2B_R by 69 ± 3.2% of PBS control group (Fig. 4E). This effect was eliminated after siRNA treatment: in the presence of BzATP the level of 5-HT_2B_R decreased, insignificantly, to 91 ± 7.3% (n = 6) of PBS group (Fig. 4E). After over-expressing FoxO3a, the basic level of 5-HT_2B_R was elevated by 62 ± 4.7% as compared with PBS group without over-expression, but BzATP did not change the expression of 5-HT_2B_R (Fig. 4E).

### 3.4 The Effects of P2X_7_R on the Phosphorylation of STAT3 and cPLA2 are Related to 5-HT_2B_R in vivo and *in vitro*

Exposure of mice to SD decreased the phosphorylation of STAT3 by 66 ± 5.4% of control, however, this effect was abolished in P2X_7_R-KO mice, the level of p-AKT in SD group was 97 ± 8.9% control (Fig. 5A). In contrast, SD had an opposite effect on the activation of cPLA2. As shown in Fig. 5B, the phosphorylation of cPLA2 in animals exposed to SD increased by 67 ± 8.7% (n = 6) as compared with controls. This effect was absent in P2X_7_R-KO mice in which SD did not stimulate the activation of cPLA2 (Fig. 5B).

**Figure 5.**
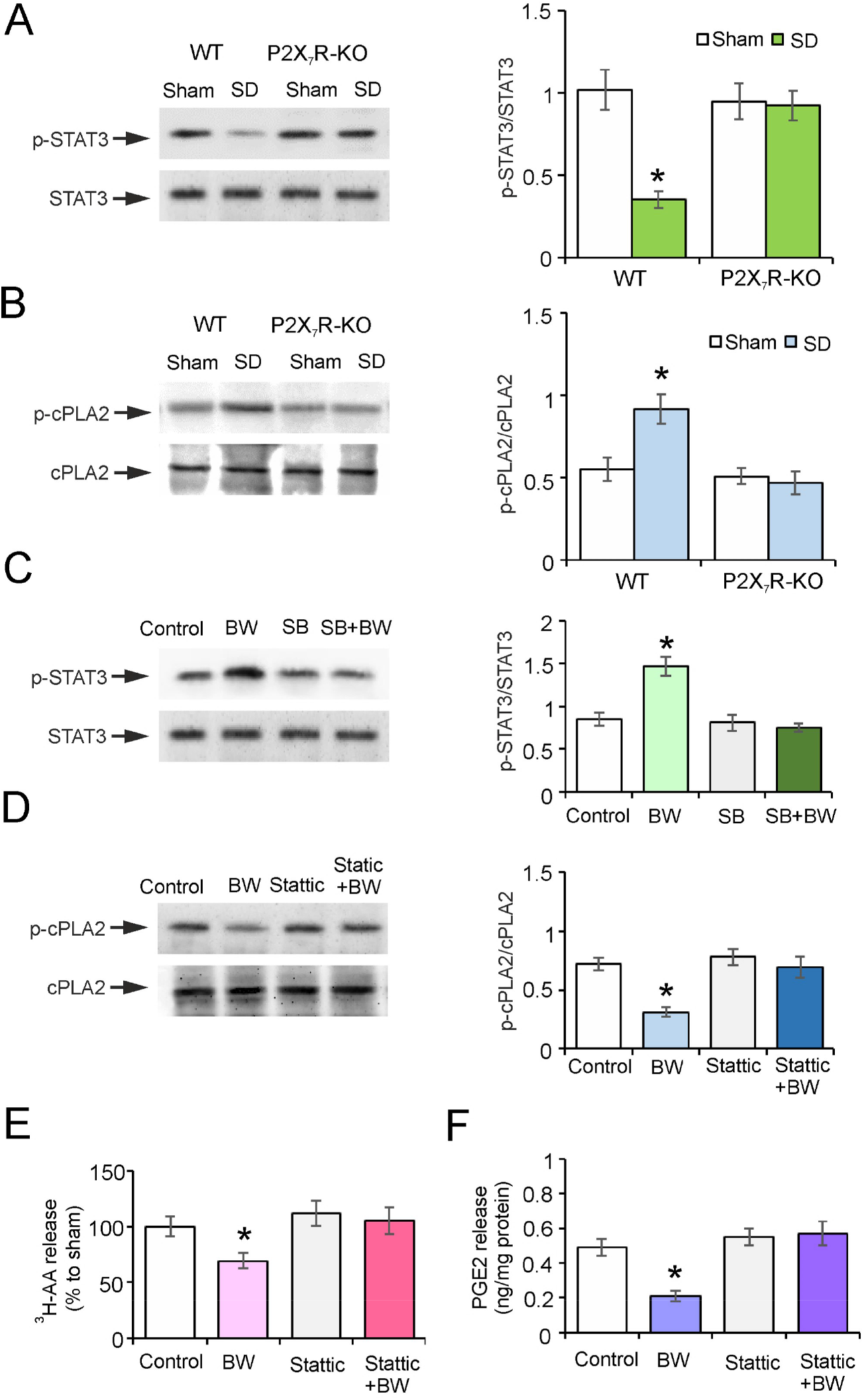
P2X7 receptors regulate 5-HT_2B_Rs-dependent activation of STAT3 and cPLA2. (A and B) The wild type (WT) and P2X7R-KO mice were treated with sham (Control) or exposed to SD for 3 weeks, the ratio of p-STAT3 and STAT3 (A2) and the ratio of p-cPLA2 and cPLA2 (B2) were analyzed. (C-F) The primary cultured astrocytes were pretreated with SB204741 (selective 5-HT_2B_R antagonist) or Stattic (STAT3 inhibitor) for 30 min, then the cells were treated with specific 5-HT_2B_R agonist BW723C86 (BW) for 1 hour, the ratio of p-STAT3 and STAT3 (C2), the ratio of p-cPLA2 and cPLA2 (D2), and the release of ^3^H-AA (E) or PGE2 (F) were prospectively measured. Data represent mean ± SEM. *Indicates statistically significant (p<0.05) difference from any other group, n = 6.

In experiments *in vitro*, we employed 5-HT_2B_R agonist BW723C86 (BW) to stimulate 5-HT_2B_R. In cultured astrocytes, BW induced the phosphorylation of STAT3 by 73 ± 10.7% (n = 6) of control group; the 5-HT_2B_R specific antagonist SB204741 totally suppressed the phosphorylation of STAT3 stimulated by BW (Fig. 5C). Treatment with BW decreased the level of p-cPLA2 by 57 ± 3.9% (n = 6) of control group, while the irreversible STAT3 activation inhibitor Stattic elevated the phosphorylation of cPLA2 to 96 ± 8.7% (n = 6) of control group, there was no difference between control and stattic groups (Fig. 5D).

The BW decreased levels of AA and PGE2 by 31 ± 7.1% and 57 ± 3.3% respectively (n = 6) when compared with the control group, (Fig. 5E and 5F). Exposure to Stattic increased the release of AA and PGE2 reduced by BW to 105 ± 12.2% and 116 ± 12.7% of control (Fig. 5E and 5F).

### 3.5 Effects of P2X_7_R on the Depressive-Like Behaviours Induced by SD

We monitored depressive-like behaviours in P2X_7_R-KO mice, as shown in Fig. 6. There was no obvious difference in body weight between the control and SD groups of wild type and P2X_7_R-KO mice (Fig. 6A). In sucrose preference test, treatment with SD decreased the uptake percentage of sucrose water by 47 ± 5.3% (n = 6) as compared with control group in WT mice (Fig. 6B). The anhedonia induced by SD was abolished in P2X_7_R-KO mice, the uptake of sucrose water was 89 ± 4.7% of control (Fig. 6B). In the tail suspension test (TST), the immobility time of SD group was prolonged to 155 ± 15.1% of control, while this time was cut down to 107 ± 11.2% control in P2X_7_R-KO mice (Fig. 6C). Similarly, the immobility time in forced swimming test (FST) was increased by 106 ± 11.7% of control group, the time was increased only to 113 ± 9.5% of sham group in P2X_7_R-KO mice (Fig. 6D).

**Figure 6.**
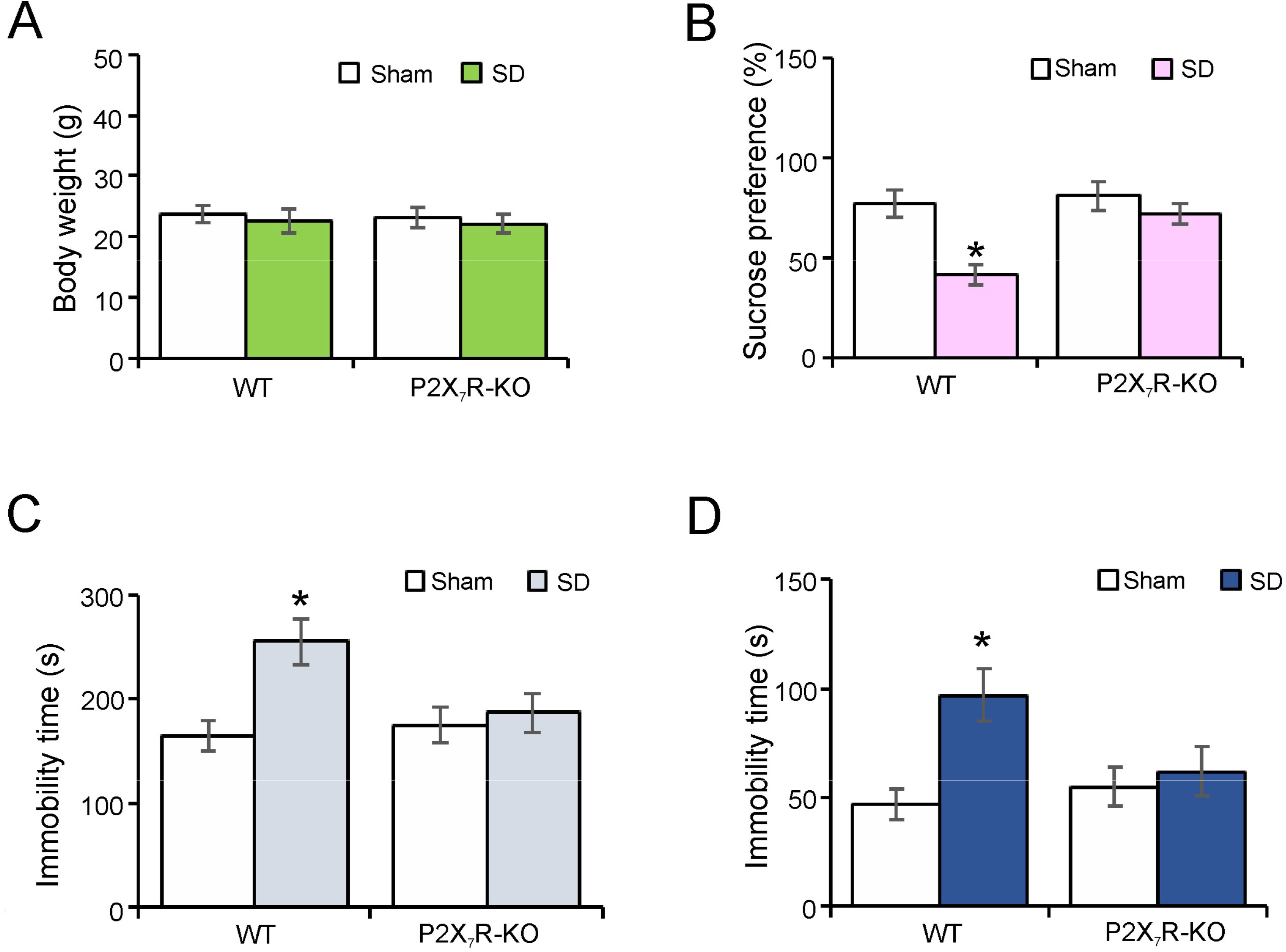
The depressive-like behaviours induced by SD are eliminated in P2X_7_R-KO mice. (A-D) The wild type (WT) and P2X_7_R-KO mice were treated with sham (Control) or exposed to SD for 3 weeks. The body weight (A) and the percentage of sucrose preference was measured (B), the time of immobility were recorded in tail suspension test (C) and force-swimming test (D), the values are expressed as mean ± SEM. *Indicates statistically significant (p<0.05) difference from any other group, n = 6.

## 4. Discussion

Our study shows that the chronic SD gradually increases the release of ATP that subsequently stimulates P2X_7_Rs which decreases the expression of 5-HT_2B_R specifically in astrocytes. Such an effect was not observed in neurones. Stimulation of P2X_7_Rs triggered by SD suppressed the expression of 5-HT_2B_R through inhibiting the phosphorylation of AKT (Ser473) and FoxO3a (Ser253) in astrocytes. The dephosphorylated FoxO3a translocates into nucleus (Li et al., 2017; Polter et al., 2009), increased FoxO3a in nucleus down-regulates the expression of 5-HT_2B_R in astrocytes. The downregulation of 5-HT_2B_R induced by SD caused the decrease in the activation of STAT3 which inhibits activation of cPLA2. As a result, chronic SD indirectly stimulated the phosphorylation of cPLA2 via down-regulating the expression of 5-HT_2B_R in astrocytes. This increased activation of cPLA2 stimulates the release of AA and PGE2, which may be linked to the depressive-like behaviours (Fig. 7).

**Figure 7.**
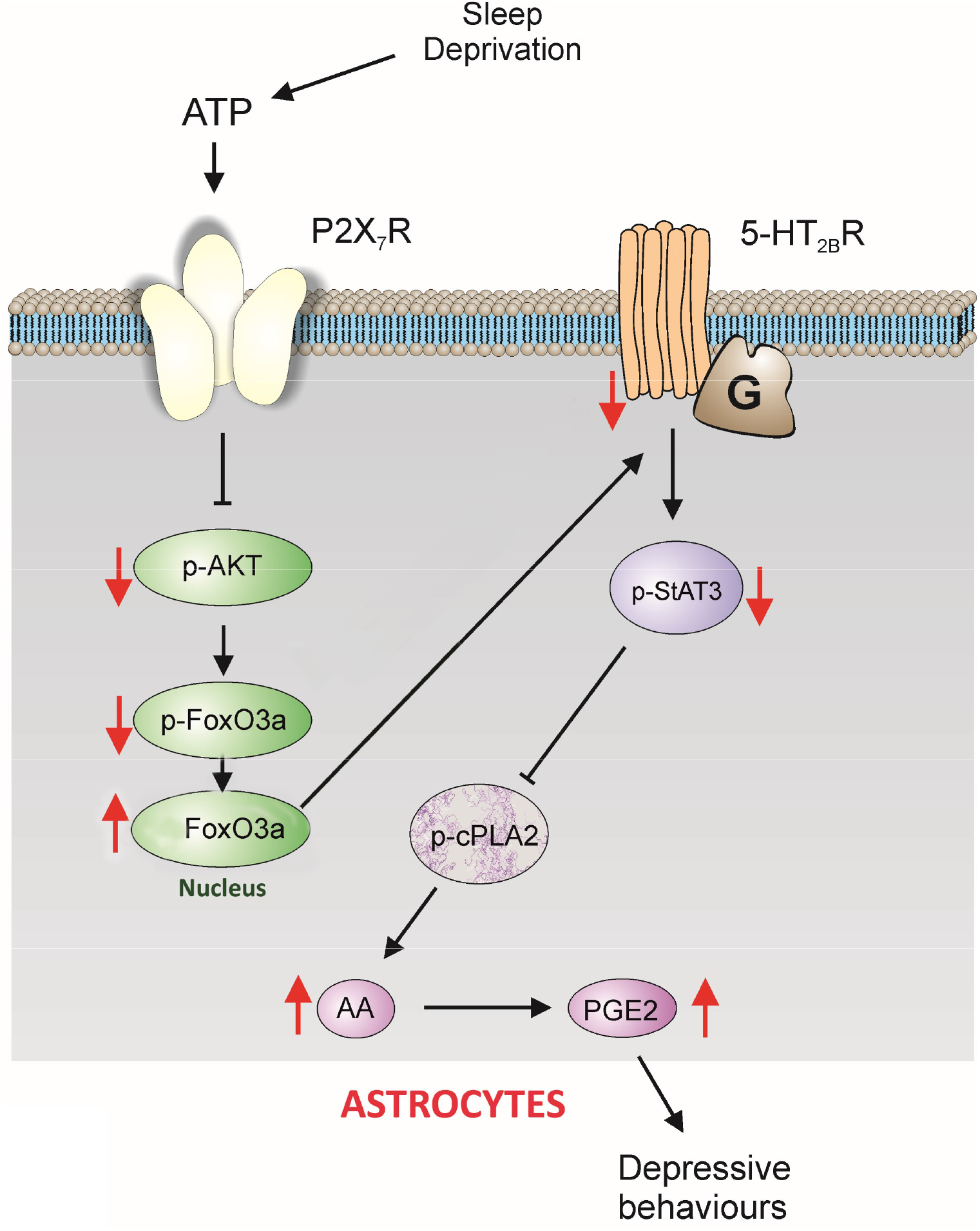
Expression and function of 5-HT_2B_ receptors are selectively decreased by SD through P2X_7_ receptors in astrocytes. The long-term treatment with SD stimulated P2X7R via ATP, the activated P2X_7_R suppressed the phosphorylation of AKT and FoxO3a in cytoplasm, the dephosphorylated FoxO3a accumulated into nucleus in astrocytes. The increased FoxO3a down-regulated the expression of 5-HT_2B_R, and the phosphorylation of STAT3 was also decreased, which relieved the inhibition on the phosphorylation of cPLA2. The activated cPLA2 promoted the release of AA and PGE2, even caused the depressive-like behaviours.

Mechanisms of sleep are complex and sleep impairments have numerous negative impacts. The sleep regulatory signal triggered by extracellular ATP activates glial P2X_7_Rs, which stimulates the release of pro-inflammatory cytokines, such as TNF-α and IL-1β (Krueger, Huang, Rector, & Buysse, 2013). Although there are some reports about the pattern of ATP dynamics in sleep or SD (Chennaoui et al., 2017; Dworak et al., 2010), fluctuation of ATP in long-term SD remained unexplored. Here we demonstrate the gradual increase in ATP levels in frontal cortex in the course of chronic SD. Elevated expression of P2X_7_R in monocytes of healthy subjects was reported to be related with a stress response to SD (Backlund et al., 2012). In addition, we found that the activation of NLRP3 inflammasomes induced by long-term SD was abolished in P2X_7_R-KO mice, the effects of long-term SD on the neuronal apoptosis was also eliminated through inhibiting the inflammasomes activation (Xia et al., 2018).

According to our previous reports, astroglial 5-HT_2B_Rs are the key target of selective serotonin reuptake inhibitors (SSRIs). In particular fluoxetine directly stimulates astroglial 5-HT_2B_R thus mediating anti-depressant action (Li et al., 2009; Li et al., 2008; Peng et al., 2014; Peng et al., 2018). Expression of 5-HT_2B_R elevated by leptin enhances positive effects of fluoxetine on the depressive-like behaviours induced by chronic SD (Li et al., 2018). In the present study, long-term SD selectively blocked the expression of 5-HT_2B_R by activation of P2X_7_R in astrocytes (Fig. 1 and Fig. 2). Activation of P2X_7_Rs induced by SD decreased the phosphorylation of AKT selectively in astrocytes, but did not change the level of p-AKT in neurones (Fig. 3A and 3C). Likewise, BzATP induced the dephosphorylation of AKT in primary cultured astrocytes (Fig. 4C), similar effects of BzATP on the dephosphorylation of AKT was also reported in microglia (He, Taylor, Fourgeaud, & Bhattacharya, 2017). At the same time BzATP did not change the activation of AKT in granule neurones (Ortega, Pérez-Sen, Delicado, & Miras-Portugal, 2009). Activated AKT phosphorylates FoxO3a at Ser253 in cytoplasm (Li et al., 2017; Polter et al., 2009). In contrast, dephosphorylation of FOXO3a causes its translocation from the cytoplasm into the nucleus, which triggers downstream gene expression (Li et al., 2017; Polter et al., 2009). We demonstrate that stimulation of astrocytic P2X_7_Rs suppressed the phosphorylation of FoxO3a *in vivo* and *in vitro* (Fig. 3B, D and 4D), which promoted the translocation of FoxO3a to the nucleus and inhibited the expression of 5-HT_2B_R. Over-expression of FoxO3a in the nucleus decreased the level of 5-HT_2B_Rs (Fig. 4E), while the FoxO3a-KO mice present obvious anti-depressive-like behaviours (Polter et al., 2009).

We previously reported that chronic SD decreases the phosphorylation of STAT3 in astrocytes (Li et al., 2018). In this study, we show that the decrease of p-STAT3 induced by SD was caused by the reduced expression of 5-HT_2B_R, because the activation of 5-HT_2B_R by an agonist (BW) increased the phosphorylation of STAT3 in astrocytes (Fig. 5C). In spinal cord astrocytes, treatment with ATP increases the activation of cPLA2, however, the pre-treatment with leptin eliminates the phosphorylation of cPLA2 induced by ATP via increasing the level of p-STAT3 and caveolin-1, which in turn reduces the release of AA and PGE2 (Li et al., 2016). cPLA2 selectively acts on AA containing acyl chains *in vitro* (Sundler, Winstedt, & Wijkander, 1994) and cPLA2 is a crucial enzyme in AA-derived eicosanoid production (Dennis, Cao, Hsu, Magrioti, & Kokotos, 2011).

PGE2 is metabolised from AA by the cyclooxygenase (COX) and it is an important regulator of chronic inflammation (Kalinski, 2012). The long-term treatment with SD induced the phosphorylation of cPLA2 and increased the release of AA and PGE2 via regulating 5-HT_2B_R in astrocytes (Fig. 5).

Both cPLA2 and COX-2 have been reported to associate with major depressive disorder (Gałecki, Florkowski, Bieńkiewicz, & Szemraj, 2010; Pae et al., 2004), the increased level of PGE2 is measured in depression, and COX-2 inhibitors are suggested to be used as antidepressants (Mueller, 2010). In this study, chronic SD induced the activation of cPLA2 which could trigger production of AA and PGE2 via P2X_7_R (Fig. 5), while the depressive-like behaviours induced by long-term SD were abolished in P2X_7_R-KO mice (Fig. 6).

In summary, our study revealed the mechanism underlying depressive-like behaviours induced by chronic SD, and discovered, for the first time, that decreased expression of 5-HT_2B_R induced by SD was mediated through P2X_7_Rs. Down-regulated 5-HT_2B_R dephosphorylated STAT3 thus relieving the inhibitory effect of STAT3 on the activation of cPLA2. Our results propose the selective agonist of 5-HT_2B_R or the reagent which could up-regulate the expression of 5-HT_2B_R may be considered as the therapeutic agents for preventing depression triggered by sleep disorders.

## Acknowledgments

This study was supported by Grant No. 81871852 to BL from the National Natural Science Foundation of China, Grant No. XLYC1807137 to BL from LiaoNing Revitalization Talents Program, Grant No. 20151098 to BL from the Scientific Research Foundation for Overseas Scholars of the Education Ministry of China, Grant No. 81200935 to MX from the National Natural Science Foundation of China, and Grant No. 20170541030 to MX from the Natural Science Foundation of Liaoning Province. Grant No. 81671862 and No. 81871529 to DG from the National Natural Science Foundation of China.

## Author Contributions

MX, DG and BL designed the experiments; MX, XL and ZL built the animal models; MX and SL operated cell culture; ZL and SL performed other experiments; MX and SSL analyzed the data; AV and BL wrote the paper.

## Data Sharing

The data that support the findings of this study are available from the corresponding author upon reasonable request.

## Abbreviations

SD: sleep deprivation;
MDD: major depressive disorder;
NLRP3: nucleotide-binding domain and leucine-rich repeat protein-3;
P2X_7_R: P2X_7_ receptors;
5-HT_2B_R: 5-HT_2B_ receptors;
FoxO3a: Forkhead box O3a;
cPLA2: Ca^2+^-dependent phospholipase A2;
AA: arachidonic acid;
PGE2: prostaglandin E2;
ATP: adenosine-tri-phosphate;
SSRIs: serotonin-specific re-uptake inhibitors;
5-HT: 5-hydroxytryptamine;
EGFR: epidermal growth factor receptor;
TCA: trichloroacetic acid;
DMEM: Dulbecco’s modified Eagle’s medium;
dBcAMP: dibutyryl cyclic AMP;
PFA: paraformaldehyde;
GAPDH: glyceraldehyde 3-phosphate dehydrogenase;
GFP: green fluorescent protein;
TST: Tail suspension test;
FST: Forced swimming test;
COX: cyclooxygenase;
WT: wild type

